# Children cooperate more successfully with non-kin than with siblings

**DOI:** 10.1101/2020.05.22.110387

**Authors:** Gladys Barragan-Jason, Maxime Cauchoix, Anne Regnier, Marie Bourjade, Astrid Hopfensitz, Alexis S. Chaine

**Affiliations:** Station d’Ecologie Théorique et Expérimentale du CNRS (UMR5321), 09200, Moulis, France; CLLE, Université de Toulouse, CNRS, 31000, Toulouse, France; Toulouse School of Economics, Institute for Advanced Studies in Toulouse, 31000, Toulouse, France

**Keywords:** Cooperation, Reciprocity, Kin selection, Evolution, Children

## Abstract

Cooperation plays a key role in advanced societies with human cooperation among kin being more prominent than cooperation among non-kin. However, little is known about the developmental roots of kin and non-kin cooperation in humans. Here, we show for the first time that children cooperated less successfully with siblings than with non-kin children, whether or not non-kin partners were friends. Furthermore, children with larger social networks cooperated better and the perception of friendship among non-friends improved after cooperating. These results indicate that non-kin cooperation in humans has deep developmental foundations which might serve to forge and extend non-kin social relationships during middle childhood and create opportunities for future collaboration beyond kin. Our results provide a new framework for future studies focusing on how and why cooperation with different classes of partners may change during development in humans as well as other long-lived organisms.

---

Cooperation is thought to have played a key role in the evolution of advanced societies, especially in humans^1–7^. Both kin-based interactions and reciprocity can promote cooperation^2–6^. Cooperation among kin can mean that benefits to the recipient of help can lead to indirect genetic benefits to the donor, an evolutionary process called “kin-selection”^8^, as described in social insects^9,10^ and a number of vertebrates^11,12^ Alternatively (or additionally), unrelated individuals who interact repeatedly, recognize each other, and remember their interactions can *reciprocate* leading to benefits to each individual in the partnership across time^13,14^ such as egg trading in fish^15,16^ and allogrooming in primates^17^. In a few cases, both mechanisms might operate in tandem. For example, food sharing in vampire bats is clearly maintained by reciprocity even if such sharing can occur among kin adding further indirect benefits^18–20^. Likewise, cooperation in humans occurs both among kin^21,22^ as well as through reciprocal interactions^6,23–27^, although human adults clearly favor kin over strangers^28–31^.

Studying the developmental foundations of kin and non-kin cooperation is critical to our understanding of the evolution and function of cooperation in longer-lived organisms and their role in the development and evolution of advanced societies^7,32^, yet appropriate tests in humans are still lacking. Among the limited number of experimental studies looking at cooperation or prosociality in children, reciprocation seems to be important^33–36^ but children tend to favor sharing with their relatives compared to strangers^37,38^ like adults. However, previous experimental studies of kin-sharing in children use third party tasks, where subjects are asked about abstract scenarios, which often give different results from direct participation in a task^39,40^ since it reflects what the participant thinks another person should do but not necessarily how they would actually behave in a real cooperative situation. In addition, third party tasks are thought to be “removed from the evolutionary mechanisms that […] likely shape these phenomena in early ontogeny” and do not “reflect the effects of the collaborative foraging context of early humans, in which one shares the spoils […] among those who took part in the *collaborative effort”*^40^. Because no experimental study has directly evaluated kin cooperation in situations in which individuals are asked to actively cooperate in a realistic setting, little is known about the developmental roots of kin cooperation in humans.

Here, we used a direct-action cooperation task to evaluate if children cooperate more with kin, friends, or non-friends. We measured cooperation using a rope pulling task (Fig. 1A) in which two children coordinate in pulling a single rope to reach a reward and succeed only if both rope ends are pulled at the same time and speed^41–43^. This task is complex and needs the active engagement of children in the task since it requires paying attention to the partners’ actions to succeed. While a previous experiment using this design began by giving the children a demonstration, we decided to render the task more difficult by omitting the demonstration and merely telling the children that they would have to work together to complete the task^43^. To examine the roles of kinship and friendship on performance by children of a cooperative task, we assigned each child a partner who could be classified as kin (a sibling), a socially close non-kin (a “best friend,” or a socially distant non-kin). Pairs were assigned by the experimenter based on questionnaires administered to teachers and children before the task and each pair was allowed to conduct the task until successful or up to three attempts if unsuccessful. In so doing, we investigated whether success in a cooperative task was linked to the degree of relatedness between partners in children.

**Fig. 1.**
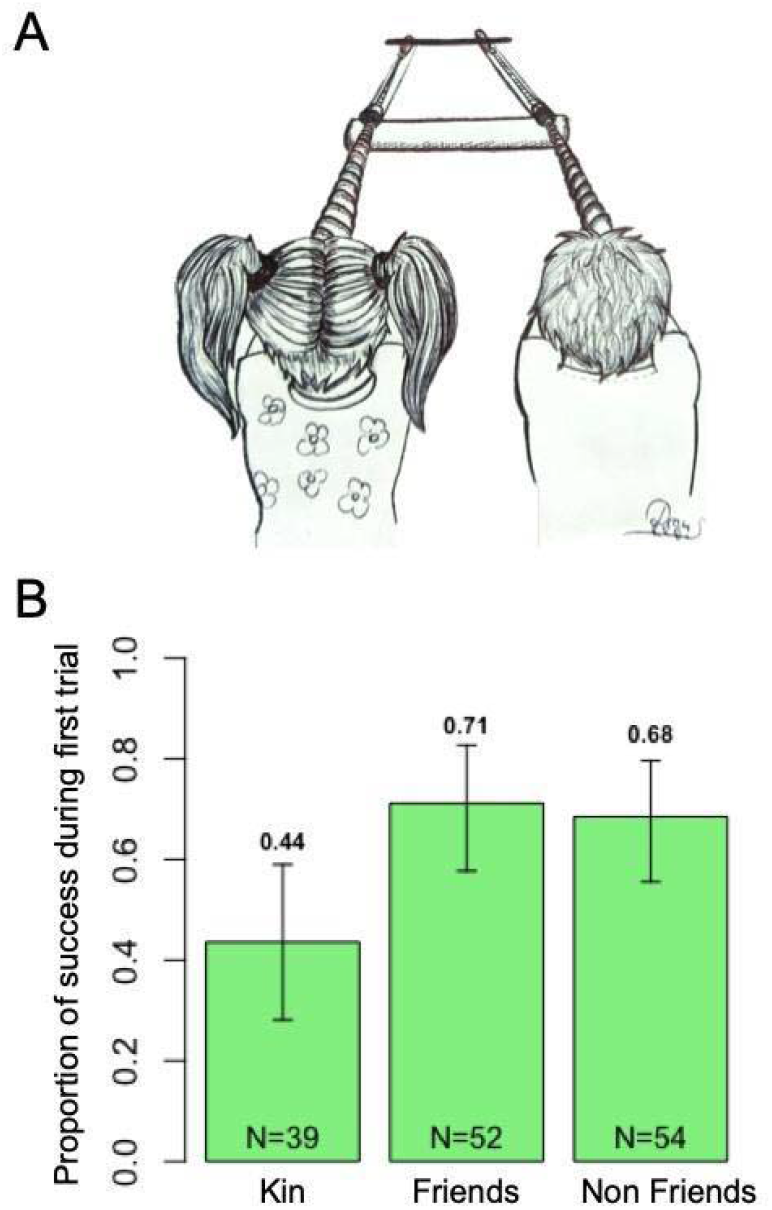
Performance in the cooperative rope pulling task by partner status. A) Illustration of the cooperation apparatus. The “rope pulling game” was adapted from previous studies on chimpanzees and children^43,45^. Photograph of the apparatus is also provided in Fig. S1 in supplemental information (SI). B) Proportion of successful first trials of kin (i.e. siblings) and non-kin friend and non-friend dyads in cooperative rope pulling. Error bars indicate 95% bootstrapped confidence interval. N values indicate the number of dyads in each category (i.e. kin, friends, non-friends).

Overall, 61% of pairs succeeded in the rope pulling task with 131 successful trials over 215 total trials performed by 145 different pairs of 290 children. Sixty-three percent (91/145) of the pairs succeeded in the first trial, 21% (31/145) in the second trial, 6% (9/145) during the third trial, and 10 % (14/145) of dyads did not succeed by the 3 trial. Overall performance in the rope pulling task was affected by the age (F=8.02; P=0.006) of the participant but not by other socio-demographic variables (binomial GLMM on overall performance including age, parents’ income, number of siblings, sex and rural vs. urban environment as fixed effects and Participant and Dyad identity as random effects, see supplementary information; supplementary methods, Table S1 and Table S2).

With direct interaction in a challenging task, we found that children readily cooperate with all categories of partners just like adults. However, contrary to what was observed in adults^21,22,44^, we found that children cooperate less well with kin than with non-kin (Fig. 1B). Kin dyads required more trials to succeed on average compared to non-kin dyads (ordered LM: friends vs. kin: t=-2.17; P=0.029 and non-friends vs. kin: t=-2.22; P=0.026) while controlling for mean age, age difference and sex (Male-Male, Female-Female and Female-Male) of dyads (ordered LM: mean age: t=-3.38; P=0.000; age difference: t=-1.21; P=0.22; sex: t=1.20; P=0.23; Table S3 in SI). Similarly, the likelihood of cooperating during the very first trial was also significantly affected by dyad type while controlling for mean age, age difference and sex of dyads (binomial GLM, partner status: deviance=7.95; P=0.019, mean age: deviance=7.29; P=0.007; age difference: deviance=2.99; P=0.084, sex: deviance=0.43; P=0.81; Fig. 1B and Table S4) with friends (52 dyads, 71% successful; z=2.62; P=0.008) and non-friends (54 dyads, 68 % successful, z=2.29; P=0.021) more likely to succeed on their first cooperative trial than kin partners (39 dyads, 44%) whereas performance of friends and non-friends did not differ from each other (z=-0.61; P=0.54; Fig. 1B).

The contrast in the propensity to cooperate with kin relative to non-kin between studies in adults and our results in children suggest that there is a striking developmental shift in the value of different forms of interactions. While we did find an improvement in success in the cooperative task with age, this improvement did not alter patterns of cooperation between kin versus non-kin while controlling for dyad age suggesting that contrasts between children and adults in their preference for cooperating with kin are not simply due to an improvement of solving a cooperative task.

More likely, the value of cooperating with different classes of individuals shifts with age. Kin cooperation among adults might provide the greatest direct and indirect benefits to success (e.g., fitness, wealth, etc.^46^) since they have reached reproductive maturity where gene transmission is likely more important than reciprocity thereby favoring kin interactions and indirect genetic benefits from cooperation. On the other hand, children are far from reproductive age and therefore might invest primarily in resource acquisition and survival which can benefit from reciprocal cooperation with peers regardless of kinship. Furthermore, kin-competition might reduce the value of kin-cooperation among children (e.g. siblings)^47–49^ especially if resources are primarily provided by parents^40^. The benefits of a given cooperative interaction to success in children are indeed modest given that children are still supported by their parents and instead may serve primarily to develop cooperative skills needed for the future such as building a social network.

Developing friendships and affiliations with peers in mid-childhood has indeed been linked to future success at adulthood^46,50,51^. Since the current network of young children is still fairly limited, reinforcing and increasing reputation through reciprocity and building a broader social network might thus be more important during childhood than adulthood. Indeed, social networks tend to expand in size among young adults, but shrink in older adults^52^. Here, we found that having a bigger social network before the experiment was related to subsequent performance during the first trial in the rope pulling task (Fig. 2) after controlling for the age difference between partners, mean age, sex and number of children in the classroom (binomial GLM, out degree centrality or number of friends named by participants: deviance=5.61; P=0.018, mean age: deviance=10.27; P=0.001; age difference: deviance=3.14; P=0.076, sex: deviance=2.12; P=0.35; number of children in the classroom: deviance=1.12; P=0.29; Fig. 3, Fig. 2, Table S5 and Fig. S3). This correlative relationship could exist either because social individuals cooperate more readily or because those who have built a bigger network develop cooperative skills. Regardless of directionality, our results show a cooperative benefit to a larger social network.

**Fig. 2.**
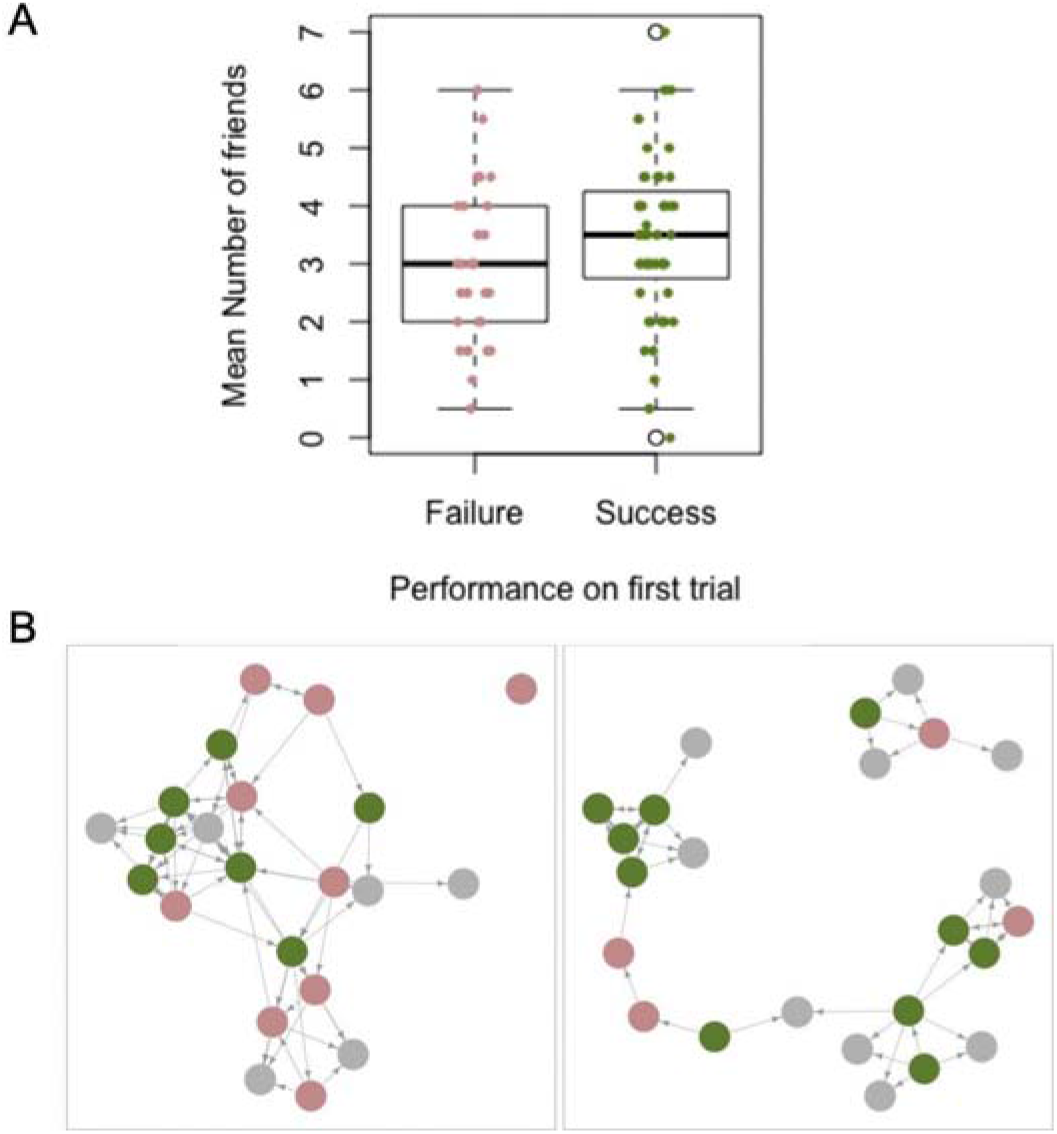
Relationship between the number of friends (out degree centrality) and performance in the rope pulling task. The number of friends and social network were based on a questionnaire before the experiment where children were asked to give the names of children they prefer to play with in their classroom including children who did not participate in the task. A) Boxplots contrasting the number of friends averaged between the two partners according to their performance during the first trial. Failure in the first trial is shown in red and success in green. Each dot represents a dyad of children. B) Examples of two classroom networks in which individuals who succeeded in the first trial appear in green and who failed in red. Children who did not participate and who participants named as friends appear in grey. Arrows represent friendship between children such that bi-directional arrows represent pairs of individuals who each listed the other as a friend whereas single headed arrows represent cases where one individual considered the other a friend while the second individual did not list the first as a friend. All networks are presented in Fig. S2.

**Fig. 3.**
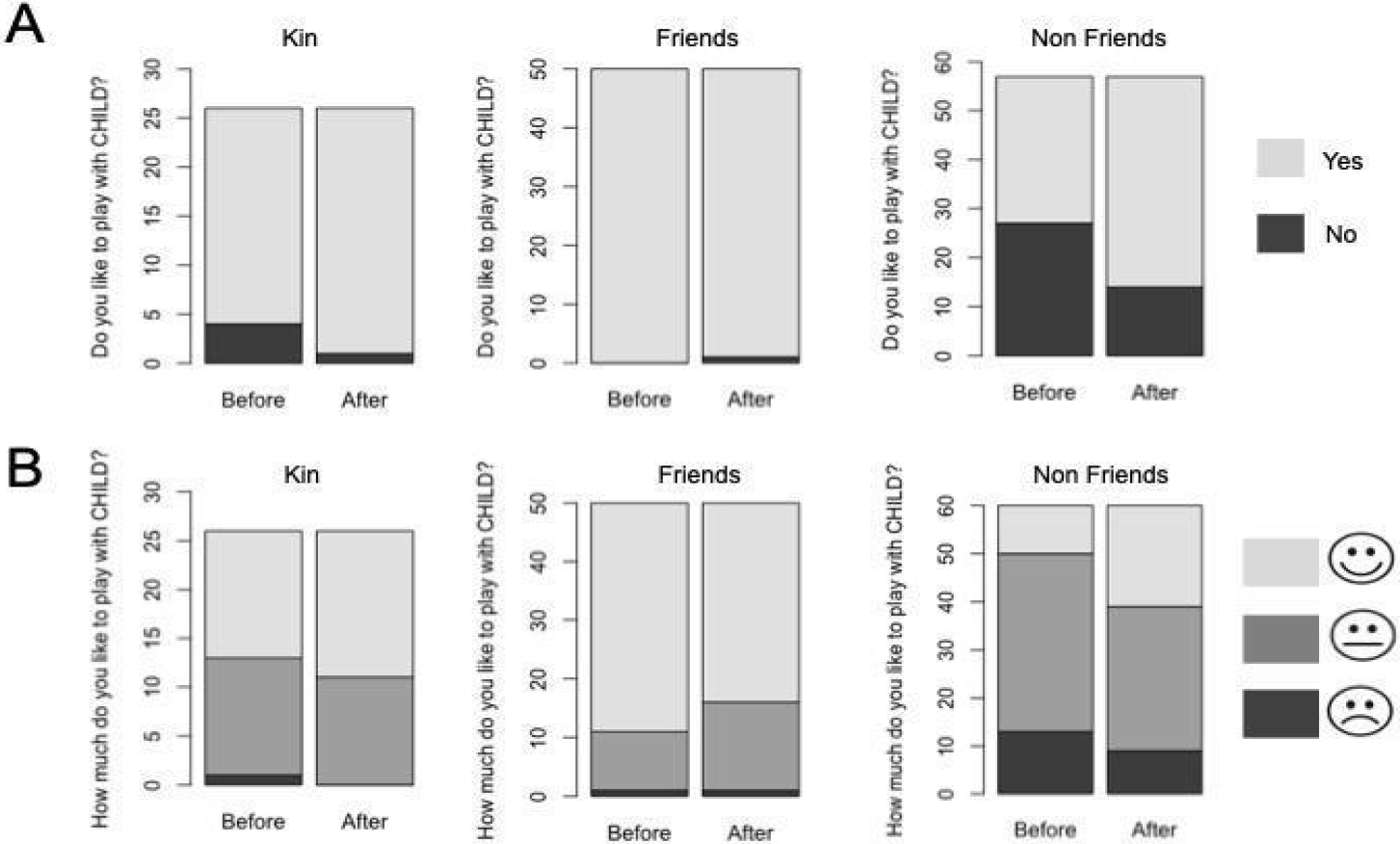
Effect of the cooperative task on the relationship quality between kin, friend and non-friend partners. A) Results from the Yes/No preference scale in which children were asked to answer the following question, before and after the rope pulling task: “Do you like to play with CHILD X?”. “Yes” responses appear in light grey and “No” responses in dark grey. B) Responses from the Emoticon Likert scale during which children were asked to rate how much they like to play with their partner twice (before and after the rope pulling task). They can either respond “A lot” in light grey, “a little” in medium grey or “not at all” in dark grey while pointing a “smiley scale” corresponding to each level.

Building a large social network should be especially valuable in unpredictable environments, since extending one’s social network to cooperate with non-kin could provide benefits when the social community is perturbed whereas limiting one’s social network only to kin would be risky^18,19^. For example, under unpredictable situations^19^ or when non-kin are more numerous than kin^18^, bats tend to favor cooperation with non-kin compared to kin partners^19^. For children, the school environment is constituted mostly of non-kin and has some risks (e.g. victimization by peers^53–55^) so expansion of the social network could indeed carry “social bet hedging” benefits. If cooperation serves to strengthen or build a social network, we would predict that participation in a cooperative action should alter future interactions. As such, we investigated whether performing the rope pulling task subsequently modified the relationship between the two partners. To do so, children were asked to rate their relationship with their partner before and again one day after the cooperative task (Fig. 3) using a Yes/No preference test (“Do you like to play with CHILD X?”; Fig. 3A) and an emoticon Likert scale (“How much do you like to play with CHILD X? A lot, a little, not at all”; Fig. 3B). While there was no change in how much children liked kin partners (Yes/No preference test: mcNemar chi2=(1,25)=1.33; P=0.25; emoticon Likert scale: Cochran Q(1,25)=1.80; P= 0.18; Fig. 4) and friend partners (Yes/No preference test: mcNemar chi2(1,49)=1; P=1; emoticon Likert scale: Cochran Q(1,49)=2.78; P=0.10; Fig. 3) after the task, we found that the relationship quality of non-friends improved after performing the rope pulling task together in both the Yes/No preference scale (mcNemar chi2(1,56)=9.60; P = 0.002; Fig. 3A) and the Emoticon Likert scale (Cochran Q(1,55)=7.35; P=0.007; Fig. 3B).

Overall, our results show that cooperation between non-kin partners plays a key role during childhood which we argue serves to expand a child’s social network since non-friends had a more positive view of their partner after interacting during the cooperative task. These results contrast with two studies in which children had to choose how to allocate resources between dolls (i.e. third party tasks with fictional characters) which both showed greater apparent cooperation with kin than friends or strangers^37,56^. We believe that differences in experimental methods lead to this contrast since the use of fictional characters is more likely to elicit a response that reflects what children think they or others *should* do^39,57^ whereas direct interactions to cooperate in a face-to-face situation used in our experiments should better reflect the actual outcome of natural cooperative situations. Furthermore, direct tasks are more challenging than simple allocation tasks such that costs of cooperation could also alter decisions. A difference in the value of friendships vs. sibling relationships among children relative to adults might drive contrasts in how much effort each group put into the cooperative task. The rope pulling task used here requires continued, active attention and coordination with a partner to succeed. Indeed, the number of gazes exchanged by the dyad in the current study was a strong predictor of success and kin displayed fewer gazes than other types of dyads as expected given that kin were less successful in the task (Supplemental information; Fig. S3). Given the difficulty of the cooperative task we conducted and considering the time children in western societies spend at school, we believe our results reflect an ecologically realistic measure^58^ of cooperation in children.

Using this direct-action cooperation task, we found that children cooperate more with non-kin peers compared to siblings in direct contrast with results in adults. At what age shifts between cooperative strategies in children and adults occur remains an open question for future research. A more complete understanding of how decisions about cooperation shift through life will require both attention to the context of testing and application of similar direct-interaction tests in individuals from a broad age range. Whether such developmental shifts in cooperation are common in other organisms remains to be explored, but should exist in cases where the benefits of cooperating with different types of partners shifts through life^59^. This new hypothesis motivated by our findings challenges our understanding of cooperation and should stimulate new research into cooperation across life stages in both humans and other social organisms.

## Acknowledgments

We thank parents of participants, directors, teachers and inspectors of the schools for permission to work in schools and children who participated in the study. We thank Orlane Scelsi for her drawing of the apparatus, David Lieuré for help adapting and building the apparatus for children and Bruce Lyon and Michael Singer for helpful feedback on the manuscript. ANR-Labex TULIP (ANR-10-LABX-41) New Frontiers Grant entitled “Human Altruism Genes” and the Institute for Advanced Studies in Toulouse provided funding. This work was supported by the Laboratoire d’Excellence (LABEX) entitled TULIP (ANR-10-LABX-41) and IAST through ANR grant ANR-17-EURE-0010 (Investissements d’Avenir program).

## Author Contributions

G. B. -J., A. S. C, M.B and A.H. designed research; G. B. -J and A. R. performed research; M. C. and G. B. -J. analyzed data; G.B.-J, A.S.C and M.C. wrote the paper; M. B. and A.H. gave feedback on the paper.

## Additional Information

Supplementary Information is available for this paper. Correspondence and requests for materials should be addressed to Gladys Barragan-Jason.

## Methods

### Participants

We recruited 290 children from ages 3 to 10 (92 3-to-5 year-olds, 139 6-7 year-olds and 59 8-to-10 year-olds; 135 females) from 15 kindergartens and elementary schools in southwestern France. All parents signed an informed consent form for their children and only children who gave their verbal assent were included. Parents also filled a demographic questionnaire including parents’ income, living area (urban vs. rural), number of siblings and native language. Thirty percent of children were from middle-class backgrounds (20,000 to 30,000 euros/year) and 35 % lived in urban areas. Participants had 2.5 siblings on average: 18% were an only child, 44% had only one sibling, 20% had two siblings and 18% had more than 2 siblings. Sixty-nine percent of the children were native French and all children (except 2 children from whom the test was performed in English) were French-speaking. The same female experimenter tested children during a single video-recorded session in an available room at their schools.

### Experimental procedure

Participants performed a rope pulling task^1^. This task requires coordinated pulling to reach a reward (see Fig. 1 and Fig. S1). Two children are required to pull their end of the rope simultaneously, each holding the end of the same rope where the two ends are far enough apart that one person could not reach both ends at the same time. A single rope is threaded around an apparatus such that only if both rope ends are pulled at the same time can the containers be moved and the rewards be reached. Pulling on one end would only move the rope but not the two sliding containers which contain the rewards, making the other end of the rope unavailable to a partner. Only if both participants pulled the rope at the same time and at similar speeds, would they each obtain a reward. Two cases lead to failure in the cooperative task: asymmetric pulling led to neither participant obtaining the reward (0/0) or if both participants pulled at the same time but one let go too quickly, then just one of the two obtained the reward (0/1 or 1/0).

The Experimenter (E) explained to the children that they would play together to win a reward (stickers) but provided no further instruction. By not providing more guidance, this cooperative task was rendered more difficult than previous studies (63% here vs. 94% during first trial in a previous study^2^). E placed the two rewards (one for each child) in the apparatus (one in each container of the apparatus) under observation of the children and told them they could start to play. The partners of a dyad could be either siblings, best friends (someone they frequently play and interact with), or non friends (someone they know but do not particularly interact with). The status of the dyad (i.e., siblings, friends, non-friends) was determined before conducting the experiment by asking the children’s teachers to name the friends of the participants and specifying each participant’s best friend through a questionnaire filled out before the experiment. We asked teachers to base their estimation of relationship closeness of a dyad on the amount of time children spent together, intensity of positive interactions, and time they play with each other at school^3,4^. Based on the responses of the questionnaires, dyads were formed by E. Only reciprocal relationships were included meaning that each individual was considered the best friend or not-friend of the other. In order to check the accuracy of the teachers’ rating, E subsequently asked children about their relationship with their partner (“Do you like to play with CHILD X?”) and about the quality of such relationship using an emoticon Likert scale (“How much do you like to play with CHILD X?: a lot, a little, not at all). The order of the emoticons was counterbalanced across participants to avoid bias. In order to investigate whether participation in the rope pulling task affected the relationship of the partners, the same questions were administered 24 h after the test. Finally, we gathered information about each child’s friend network before the experiment, by asking the child to name their friends (“Please, tell me the names of the children you like to play with the most?”). Due to logistical constraints, we were only able to gather complete friendship network data at 10 schools of the 15 schools in our sample.

Each time a pair attempted to pull the ropes is termed a “trial”. Dyads of children could perform a maximum of 3 trials beyond which E stopped the testing and gave the stickers to the children. Overall, most children succeed within those 3 trials as only 14 dyads failed. Overall the sample included 39 pairs of siblings, 52 pairs of friends, and 54 pairs of non-friends who performed 77, 71 and 67 trials respectively, corresponding to a total of 215 trials among 145 dyads.

### Data coding

All trials were recorded using a video camera oriented so that both participants and the apparatus were visible allowing us to score the children’s performances. Successful trials were scored when both partners pulled together and successfully reached the reward. Failed trials were scored when children failed to pull the rope together such that neither reached the reward or when only one child reached the reward. The same trained research assistant coded the number of gazes (each movement of the eyes accompanied by a movement of the head toward the partner) of each dyad during the first trials. The number of gazes include situations when both partners look at each other and when a single individual looks at the other one.

### Statistical analysis

All analyses were performed in the R environment for statistical computing version 3.3.6 (R Development Core Team, 2018).

We first investigated whether overall performance was impacted by individuals’ socio-demographic variables using generalized linear mixed models with a binary response (1: success, 0: failure;^5^ including parents’ income (low class, middle class, high class), living area (urban vs. rural), number of siblings (“0”, “1”, “2” or more than 2), age in months and sex as fixed factors and the participant as a random factor. We also investigate the effect of these demographic variables on performance on the first trial using a binomial GLM.

We then looked at the effect of partner relationship on the number of trials before success using an ordered logistic regression (“polr” function) with ordered responses (1: success after 1 trial; 2: success after 2 trials, 3: success after 3 trials or failure). We built a full model that included fixed effects of dyad relationship (kin, friend, non-friend), average age of partners, age difference between the partners and sex of the dyad (male-male, female-female, male-female).

We then examined the effect of partner relationship within a dyad on cooperation using a binomial GLM including the first trial (0 vs. 1) performed by each dyad. We built a full model that included fixed effects of dyad relationship (kin, friend, non-friend), dyad sex (male-male, female-female, male-female), average age of partners and age difference between the partners. We also looked at the impact of gaze frequency (gazes/second) using a GLM including the first trial (0 vs. 1) performed by each dyad. We built a full model that included fixed effects of dyad relationship (kin, friend, non-friend), dyad sex (male-male, female-female, male-female), average age of partners and age difference between the partners. We also investigated the effect of partner status on the number of gazes using a LM including gaze frequency (gaze per second) during the first trial. We included dyad relationship (kin, friend, non-friend), dyad sex (male-male, female-female, male-female), average age of partners and age difference between the partners as fixed effects.

We assessed the relationship between performance in the cooperative task and the size of a child’s social network. Using a binomial GLM, we asked whether performance (0 vs. 1) during the first trial was affected by a child’s number of friends (i.e. outdegree centrality in social network analysis) while controlling for dyad sex, mean age of the dyad, age difference of the dyad partners, and number of children in the classroom.

For GLMs, visual inspection of residual plots using the DHARMa package^6^ did not reveal deviations from homoscedasticity or normality. For each fixed effect, statistical significance was evaluated by likelihood ratio tests of the full model against the same model without the tested fixed effect. We report likelihood ratio F values or deviance and P-values as well as marginal, conditional or pseudo R2 of the Full Model when appropriate.

Finally, we tested the effect of the cooperative task on the quality of the relationship between the two partners, we performed McNemar and Cochran Q test when appropriate on kin, friend and non-friend partners separately.

## Notes

### Competing Interest Statement

The authors have declared no competing interest.

