## Supplemental information for "Children cooperate more successfully with non-kin than with siblings"

**Table S1.**

Table S1. Effect of demographic variables on overall performances with “participant” and “Dyad Identity” as random factors and Age in months, sex, number of siblings, parents’ income, and living area as fixed factors using binomial GLMM.

|  |  | Df | F | P | Conditional R2 | Marginal R2 |
| --- | --- | --- | --- | --- | --- | --- |
| Overall Performance  (0 vs. 1) | Binomial GLMM | - | - | - | 0.519 | 0.07 |
|  | Age | 1 | 8.02 | **0.006** | - | - |
|  | Sex | 1 | 0.08 | 0.79 | - | - |
|  | Siblings | 3 | 0.13 | 0.72 | - | - |
|  | Income | 2 | 0.32 | 0.54 | - | - |
|  | Living area | 1 | 0.004 | 0.97 | - | - |

**Table S2.**

Table S2. Effect of demographic variables on first trials performances with “Dyad Identity” as random factor and Age in months, sex, number of siblings, parents’ income, and living area as fixed factors using binomial GLMM.

|  |  | Df | F | P | Conditional R2 | Marginal R2 |
| --- | --- | --- | --- | --- | --- | --- |
| Overall Performance  (0 vs. 1) | Binomial GLMM | - | - | - | 0.99 | 0.0016 |
|  | Age | 1 | 0.014 | 0.55 | - | - |
|  | Sex | 1 | 0.00 | 0.95 | - | - |
|  | Siblings | 3 | 0.001 | 0.86 | - | - |
|  | Income | 2 | 0.001 | 0.83 | - | - |
|  | Living area | 1 | 0.001 | 0.83 | - | - |

**Table S3.**

Table S3. Effect of dyad characteristics (partner status: Kin, Friends, Non-Friends, sex: Female-Female, Male-Male, Female-Male, mean Age, age difference) on the number of trials before success (ordered LM)

|  |  | Df | t values | P | Pseudo R2 |
| --- | --- | --- | --- | --- | --- |
| Number of trials before success | Ordered LM | - | - | - | 0.111 |
|  | Mean age | 1 | -3.383 | **0.000** | - |
|  | Age difference | 1 | -1.212 | 0.218 | - |
|  | Partner Status | 3 | - | **0.035** | - |
|  | Sex | 2 | 1.202 | 0.228 | - |

**Table S4.**

Table S4. Effect of dyad characteristics (partner status: Kin, Friends, Non-Friends, sex: Female-Female, Male-Male, Female-Male, mean Age, age difference) on performance in the first trial (binomial GLM).

|  |  | Df | Deviance | P | Marginal R2 |
| --- | --- | --- | --- | --- | --- |
| Performance on the first trial  (0 vs. 1) | Binomial GLM | - | - | - | 0.156 |
|  | Mean age | 1 | 7.290 | **0.007** | - |
|  | Age difference | 1 | 2.993 | 0.084 | - |
|  | Partner Status | 2 | 7.950 | **0.019** | - |
|  | Sex | 2 | 0.431 | 0.806 | - |

**Table S5.**

Table S5. Effect of number of friends on success during the first trial while controlling for dyad characteristics (sex: Female-Female, Male-Male, Female-Male, mean Age, age difference) on performance in the first trials (binomial GLM).

|  |  | Df | Deviance | P | Marginal R2 |
| --- | --- | --- | --- | --- | --- |
| Performance on the first trial  (0 vs. 1) | Binomial GLM | - | - | - | 0.268 |
|  | Number of friends | 1 | 5.614 | **0.018** | - |
|  | Mean age | 1 | 10.266 | **0.001** | - |
|  | Age difference | 1 | 3.139 | 0.076 | - |
|  | Sex | 2 | 2.123 | 0.346 | - |
|  | Number of children in the classroom | 2 | 1.125 | 0.289 | - |

**Figure S1.**

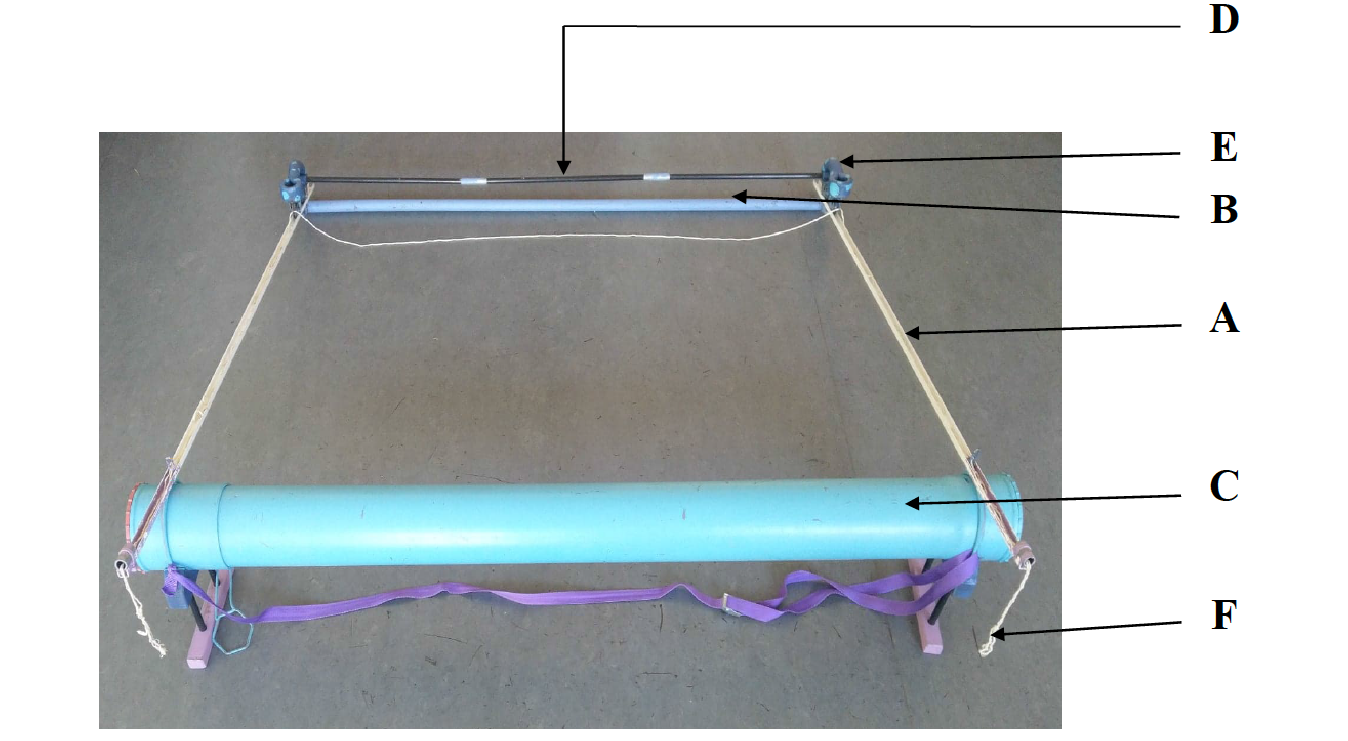

*Fig. S1. Photograph of the cooperation apparatus. The "rope pulling game" was adapted from previous studies on chimpanzees and children. It consisted of two rails (A) 150 cm long and spaced 145 cm apart attached to a metal rod (B) and a plastic tube (C). Two small metal trolleys (D) with casters were recessed on each of the rails and were connected to each other by a plastic rod (E). These two carriages had a cavity in which the reward (stickers) was placed. A 480 cm rope was threaded into different notches at the rail and trolleys in such a way that if only one of the children pulled on one end of the rope (F), the other end was automatically pulled in the rail out of reach of the second child. The rope extended the end of the apparatus by only15 cm on each side so that one child could not pull on both ends alone. At each beginning test, the carriages were positioned on the far side of the metal rod (farthest from children). In order to access the stickers, the children had to simultaneously pull at each of the two ends of the rope and thus roll the carriages towards them to the tube. The rails were inclined at an angle of 30° so that the carriages went back if one of the two children let go of the rope.*

**Figure S2.**

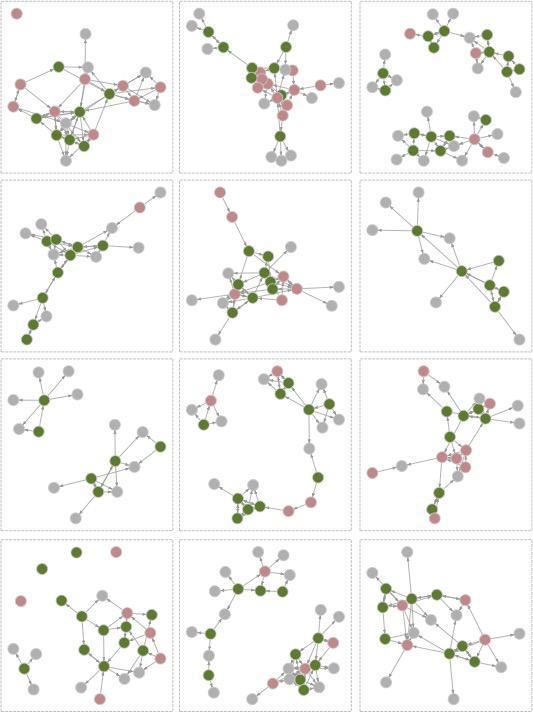

*Fig. S2. Network graphs representing relationships between children in 10 schools (12 classrooms) where data was available based on questionnaire data from children who participated in the task. Individuals who were successful in the first trial are shown in green while those who failed in the first trial are shown in red. Children who did not participate and who participants named as friends appear in grey. Arrows represent friendship between children such that bi-directional arrows represent pairs of individuals who each listed the other as a friend whereas single headed arrows represent cases where one individual considered the other a friend while the second individual did not list the first as a friend.*

**Figure S3.**

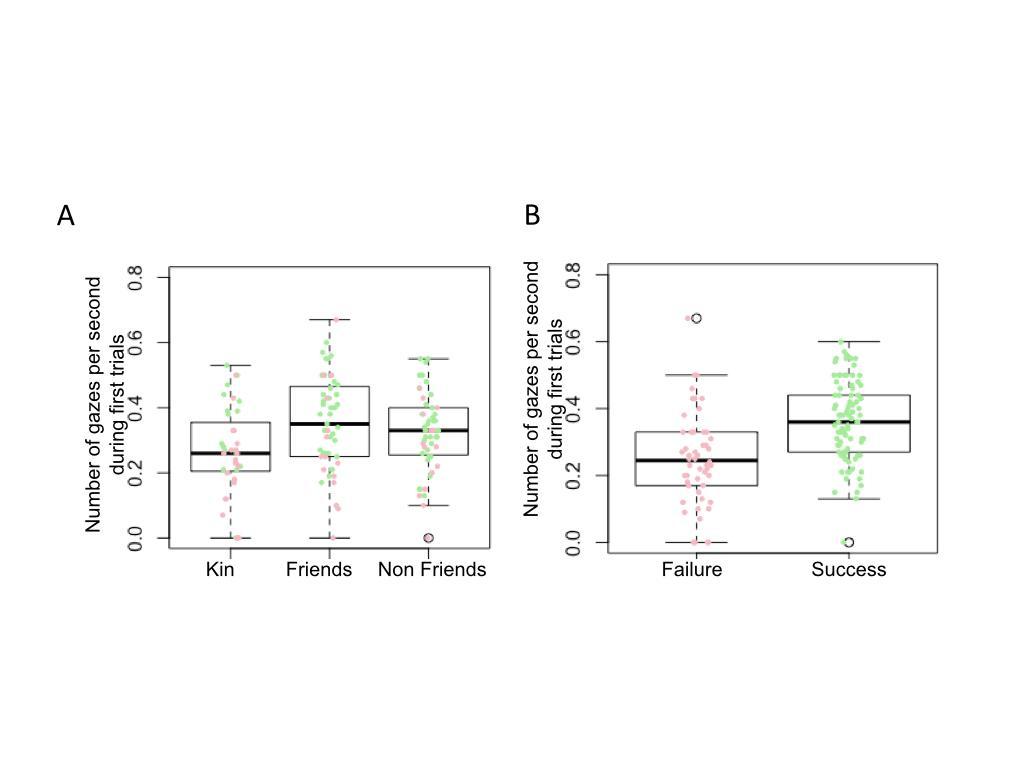

*Fig. S3. Number of gazes displayed during the first trials. A) Boxplots represent the number of gazes by second for each dyad category (i.e. Kin, friends and non-friends). Each dot represents a dyad with red dots associated with failure and green dots associated with success.*

Supplemental results

In Fig. S3, the number of gazes during the first trial was a strong predictor of success (GLM P<0.001) while controlling for the mean age (P<0.001), age difference of the dyad (P>0.05), dyad sex (P=0.053), and partner status (P=0.01). Again, kin partner displayed significantly fewer gazes than friend partners (P < 0.001) but did not differ from non-friend partners (while a tendency was observed P = 0.054), and friend partner displayed significantly more gazes than non-friends (P=0.034).
